# Uncovering potential interventions for pancreatic cancer patients via mathematical modeling

**DOI:** 10.1101/2022.01.11.475711

**Authors:** Daniel Plaugher, Boris Aguilar, David Murrugarra

**Affiliations:** Department of Mathematics, University of Kentucky, Lexington, Kentucky, USA; Institute for Systems Biology, Seattle, Washington, USA

**Keywords:** Pancreatic Cancer, Phenotype Control, Cytokines, Pancreatic Stellate Cells, Boolean Networks

## Abstract

Pancreatic Ductal Adenocarcinoma (PDAC) is widely known for its poor prognosis because it is often diagnosed when the cancer is in a later stage. We built a Boolean model to analyze the microenvironment of pancreatic cancer in order to better understand the interplay between pancreatic cancer, stellate cells, and their signaling cytokines. Specifically, we have used our model to study the impact of inducing four common mutations: KRAS, TP53, SMAD4, and CDKN2A. After implementing the various mutation combinations, we used our stochastic simulator to derive aggressiveness scores based on simulated attractor probabilities and long-term trajectory approximations. These aggression scores were then corroborated with clinical data. Moreover, we found sets of control targets that are effective among common mutations. These control sets contain nodes within both the pancreatic cancer cell and the pancreatic stellate cell, including PIP3, RAF, PIK3 and BAX in pancreatic cancer cell as well as ERK and PIK3 in the pancreatic stellate cell. Many of these nodes were found to be differentially expressed among pancreatic cancer patients in the TCGA database. Furthermore, literature suggests that many of these nodes can be targeted by drugs currently in circulation. The results herein help provide a proof of concept in the path towards personalized medicine through a means of mathematical systems biology. All data and code used for running simulations, statistical analysis, and plotting is available on a GitHub repository at https://github.com/drplaugher/PCC_Mutations.

## 1 Introduction

PDAC is well known for its grim prognosis. As such, it is the fourth highest cause of cancer-related death in the United States and seventh globally, and there is only a 3%five year survival rate among its victims [5]. Pancreatic cancer is also recognized for its resistance to most traditional therapy protocols such as chemotherapy, radiotherapy, and curative resection. PDAC is difficult to detect, partially because the pancreas is located deep within body. This means that a standard doctor’s exam will likely not reveal a tumor, in conjunction with an absence of detecting and imaging techniques for early stage tumors. Treatments are often unsuccessful because diagnoses are frequently given at late stages of the disease, leaving little time to successfully eradicate and reverse its impact [27, 34].

*In silico* models are commonly used in cancer research for the discovery of general principles and novel hypotheses that can guide the development of new treatments. It is eventually possible that, when combined with cancer data and modern control techniques, *in silico* models will predict clinically relevant endpoints and find optimal control interventions to stop cancer progression. Despite their potential, concrete examples of predictive models of cancer progression are scarce. One reason is that most models have focused on single–cell type dynamics, ignoring the interactions between cancer cells and their local microenvironment. Indeed, there have been a number of models that were used to study gene regulation at the single-cell scale, such as macrophage differentiation [7, 26, 28], T cell exhaustion [4], differentiation and plasticity of T helper cells [21, 30], and regulation of key genes in different tumor types [8, 22], including pancreatic cancers. These models are great steps towards control based treatment optimization. However, it has been demonstrated that that the local microenviroment have a critical effect on the behaviour of cancer cells [33]. Consequently, ignoring the effect of cells and signals of the local microenvironment can generate confounding conclusions.

In order to study the interplay between cancer cells and other cells of the tumor microenvironment, modelers developed multicellular models including cancer, stromal, immune, and other cells of the tumor microenvironment [2]. These models are typically multiscale integrating interactions at different scales, making it possible to simulate clinically relevant spatiotemporal scales, and at the same time simulate the effect of molecular drugs on tumor progression [11–14, 18, 25]. The high complexity of these models generates challenges for model validation such as the need to estimate too many model parameters. Moreover, even though a multi-scale model would likely provide more realistic simulations, their complexity prevents the application of state of the art control techniques to find optimum therapeutic alterations based on these models.

In this work, we used a previously developed multicellular model of pancreatic cancer based on a Boolean network approximation (Fig 1), and we used control strategies that direct the system from a diseased state to a healthy state by suppression (or expression) and disruption of specific signaling pathways. The models consists of pancreatic cancer cells, pancreatic stellate cells, cytokine molecules of diffusing in the local microenvironment, and internal gene regulations for both cell types [27]. Although the model includes several components of the tumor microenvironment, its Boolean nature allows the application of recent control techniques.

**Fig 1.**
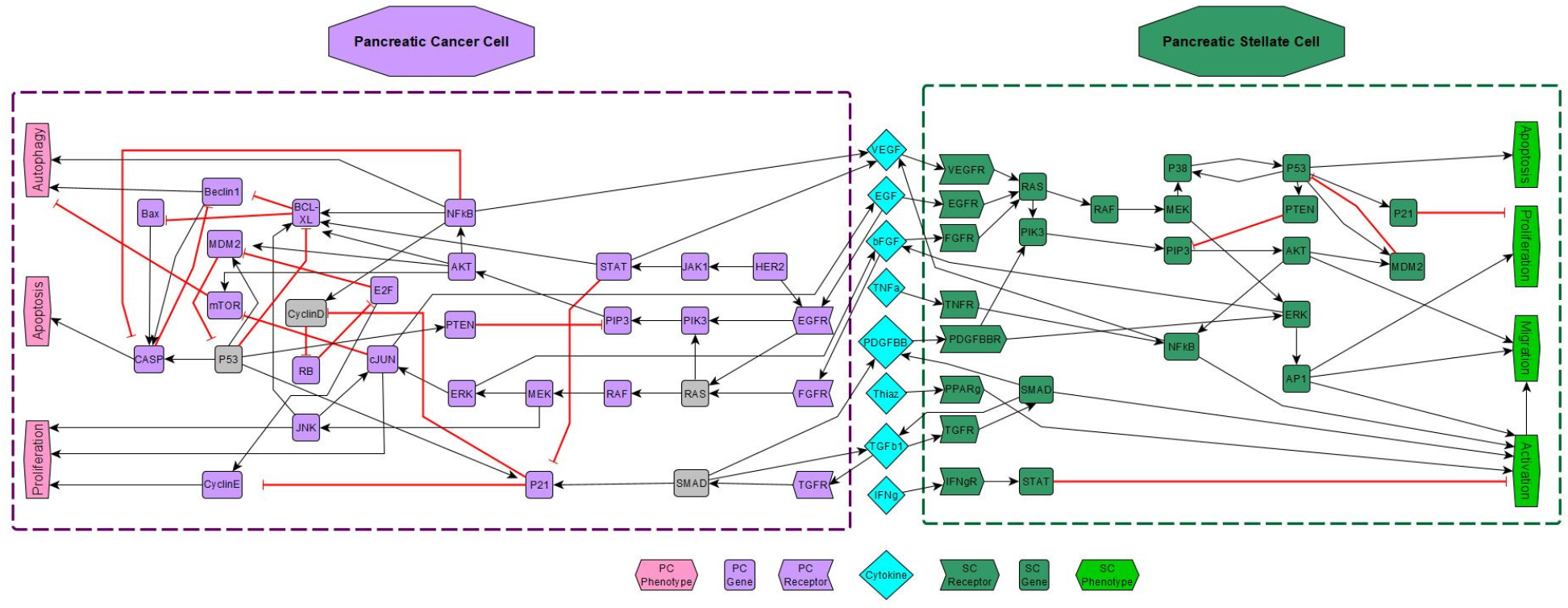
Gene regulatory network model of pancreatic cancer including PCCs, PSCs, and extracellular molecules. The purple dashed box represent PCC while the green dashed box represents PSC. Shapes and colors of nodes indicate their function and cell type (respectively), as shown in the legend. Black barbed arrows indicate signal expression, while red bar arrows indicate suppression. Grey nodes located in the PCC indicate prevalent mutant genes [27].

Further, we used techniques from phenotype control theory [24, 27, 32, 36], which is primarily concerned with identifying key markers of the system that aid in understanding the various functions of cells and their molecular mechanisms. In a biological sense, phenotypes represent the observable traits of an organism. In a similar fashion, we associate a phenotype with group of attractors where a subset of the system’s variables have a shared state. These shared states are then used as biomarkers. Phenotype control was used to identify sets of intervention targets that, if applied simultaneously to both cancer and stellate cells, induce cancer cell apoptosis.

In this paper, we used the model to study the impact of mutations commonly found in PDAC patients such as KRAS, TP53, SMAD4, and CDKN2A. We applied phenotype control theory in an effort to derive new intervention targets for particular induced mutation combinations. Control sets contained nodes within both the pancreatic cancer cell and the pancreatic stellate cell, including PIP3, RAF, PIK3 and BAX in pancreatic cancer cell as well as ERK and PIK3 pancreatic stellate cell. Throughout this writing, we will often denote PCC components with a subscript c and PSC components with subscript *s*. Within the various mutation combinations, we used our stochastic simulator based on the stochastic discrete dynamical system (SDDS) framework [23] to derive aggressiveness scores from simulated attractor probabilities as well as long-term trajectory approximations. Results showed that certain mutation combinations may indeed be more aggressive than others. We then performed statistical analysis on clinical gene expression data to identify significantly impacted genes, which corroborated targets discovered with algebraic methods. Lastly, we derived survival curves for the clinical data that served as a validation for aggressiveness scores derived by our SDDS simulator.

### 1.1 Mutation cascade analysis

PDAC has four common gene mutations: KRAS, TP53, SMAD4, and CDKN2A [2, 6, 37]. Notably, the HER2/neu oncogene mutation typically occurs in the precancerous stage, where HER2 binds to EGFR and activates its pathway. However, prevalence of the HER2 mutation is so low that we elected to exclude it from this project’s analysis [2]. Within our diagram (Fig 1), we differentiate these popular mutations in grey to easily see their signaling cascades.

Of these four mutations, the most prevalent is KRAS, occurring in nearly 95%of cases. KRAS encodes K-Ras4A or K-Ras4B, with KRAS4B being the predominant transcript [6]. KRAS oncogene mutation occurs in the precancerous stage, which then leads to continuous expression of the RAS protein. As a result, the MAPK pathway becomes overexpressed and promotes PCC proliferation through ERK and JNK activation [35]. KRAS can also upregulate the PIK3/AKT pathway, which is known promote cancer progression [9].

TP53 encodes the p53 tumor suppressor, a regulator of the cell cycle that is responsible for maintaining cellular and genetic stability [6]. This mutation typically occurs in late stages of the disease, and it induces lipid accumulation. As a result, we begin to see anticancer effects through a paracrine mechanism in the cancer microenvironment [37].

SMAD4 encodes the tumor suppressor protein Smad4, which is activated by TGFβ1, then relocates to the nucleus to begin apoptosis [35]. In the late stages of cancer, the loss of SMAD results in an increase in TGFβ1 which then promotes tumor progression through overexpression of the MAPK and PIK3/AKT pathways [3].

CDKN2A is involved in cell cycle regulation, differentiation, senescence and apoptosis through p16INK4a, p14 alternate reading frame (p14ARF), cyclin-dependent kinase4 p15 (p15INK4A) and long-chain non-coding RNA (lncRNA) ANRIL (also known as CDKN2B-AS). Missing CDKN2A prevents CDK4/6 from binding to D-cyclin, which then leads to retinoblastoma (RB) binding to E2F. The result is cell cycle arrest and cell senescence (see Fig 1). If p16INK4a is missing, then CDK4 is overexpressed. As a result, we see proliferation of B cells, rising insulin secretion and pancreatic hyperplasia. p16INK4a regulates cyclin D1 expression, and D1/CDK4 is critically involved in cellular metabolism and cell cycle progression, which provides therapeutic potential for inhibiting the progression of pancreatic cancer by cell cycle suppression [37]. In our model, we use CyclinDc as the indicator for CDKN2A due to their corresponding signaling status seen in [2].

## 2 Results

### 2.1 Finding intervention targets

To find controls for the various mutation combinations, we used the algebraic control approach in [24] with the objective of blocking undesirable phenotype expression as was done in [27]. We sought for two types of interventions: node controls and edge controls. These two strategies, though similar, provide a range of biological options. One reason is that node control requires an entire node to be knocked out (or continually expressed), but edge control simply requires an edge communication to be blocked (or continually expressed). For node control, we encode the nodes of interest as control variables, and then the control objective is described as a system of polynomial equations that is solved by computational algebra techniques. Likewise, to achieve edge control we encode the edges as control variables. This technique is further detailed in Section 4.3. Our overall objective was to find control nodes and edges that block PCC proliferation and autophagy, while upregulating PCC apoptosis. It has been hypothesised that once PSCs become activated, they begin to migrate towards PCCs to form a type of protective layer [35]. Thus, we further sought to inhibit PSC proliferation, migration, and activation, while promoting PSC apoptosis.

Among node controls discovered in the PSC, simulations indicated that knocking out ERKs suppresses all PSC phenotype expression with the exception of activation. That is, the PSCs become dormant and are not proliferating, not apoptotic, nor are they migrating. But if one wishes to induce PSC apoptosis, simulations indicated that knocking out both PIK3_s_ and ERK_s_ will increase apoptosis as well as reduce the levels of activation. Edge controls include pathways knockouts of ERK_s_→AP1_s_ along with PIK3_s_→PIP3_s_. These particular PSC controls hold regardless of mutation combination.

After inducing the KRAS mutation, we discovered node controls involving PIP3_c_ knockout combined with the previously mentioned PSC controls. Remaining mutations proved controllable by combining PIP3_c_/RAF_c_ node knockout, together with the PSC controls. Edge controls include pathways knockouts of PIP3_c_→AKT_c_, and RAF_c_→MEK_c_ in addition to the aforementioned PSC edge controls.

Interestingly, the significant genes discovered through statistical testing (MDM2_c_, PIK3_c_, and BAX_c_ - see Section 2.2) were also listed as potential targets through the algebra method. The statistical analysis discussed in subsequent sections served as a validation and filter for parsing potential targets found with algebra. Since most mutation combinations considered significant (KRAS, TP53/KRAS, TP53/SMAD/KRAS, TP53/CDKN2A/KRAS) contain TP53, MDM2_c_ only works with the KRAS mutation. This is due to a negative feedback loop involving both TP53_c_ and MDM2_c_, see Fig 1. Next, simulations confirm that PIK3_c_ works the same as PIP3_c_ due to the topology of the system. We also discovered that constant expression of BAX_c_ works best in combination with PIK3_c_ or PIP3_c_, so much so that we saw better levels of apoptosis when compared to RAFc. As with other target options, the correlating edge controls of MDM2_c_→TP53_c_, BAX_c_→CASP_c_, and PIK3_c_→PIP3_c_ proved to be as effective as the node controls because their phenotype expression levels were negligibly different.

To a great degree, the uncovering of the need for combining controls is consistent with findings in [34] because singleton controls are not always sufficient. Vundavilli et al tested the efficacy of drugs on pathways similar to those found in our model, and they showed evidence that particular drugs may work better in combination rather than individually. In the *Appendix*, Table 7 shows the simulated expressions for individual node knockout in order to verify that specific mutation combinations may require a combination of drugs.

In Table 1, we have listed the PCC targets discovered as well as potential drugs found in literature [20, 34, 35]. To demonstrate control efficacy, Figs 2 and 3 comparatively show simulation results of our node and edge controls for the key mutations studied in Section 2.2. Expression levels were determined by running 1000 random initializations for 300 time steps (with 1% noise), then recording the average expression at the last time point. See that the PSC interventions knockout all diseased PSC phenotypes except a fractional level of activation. Results from the edge control simulations are nearly the same as those with node control. However, it may be easier to block a signal rather than knockout a gene altogether. Using a combination of algebra, network topology analysis, and simulations, we uncovered intervention targets of Set 1 (PIK3_c_ / BAX_c_ / ERK_s_ / PIK3_s_) and Set 2 (PIP3_c_ / RAF_c_ / ERK_s_ / PIK3_s_). Among these particular mutation combinations, it is clear that Set 1 outperforms Set 2 in scenarios involving mutated TP53. More discussion on this phenomenon is found in Section 2.3.

**Table 1.** PCC controls with potential drugs. This table is a list of the PCC targets we discovered, with the required state in addition to potential drugs that target the given pathway.

| Target | State | Drug | Reference |
| --- | --- | --- | --- |
| PIP3 | OFF | LY294002 | [34] |
| PIK3 | OFF | LY294002 | [34] |
| BAX | ON | Venetoclax, Navitoclax | [20] |
| RAF (MAPK) | OFF | gemcitabine | [35] |
| MDM2 | OFF |  |  |

**Fig 2.**
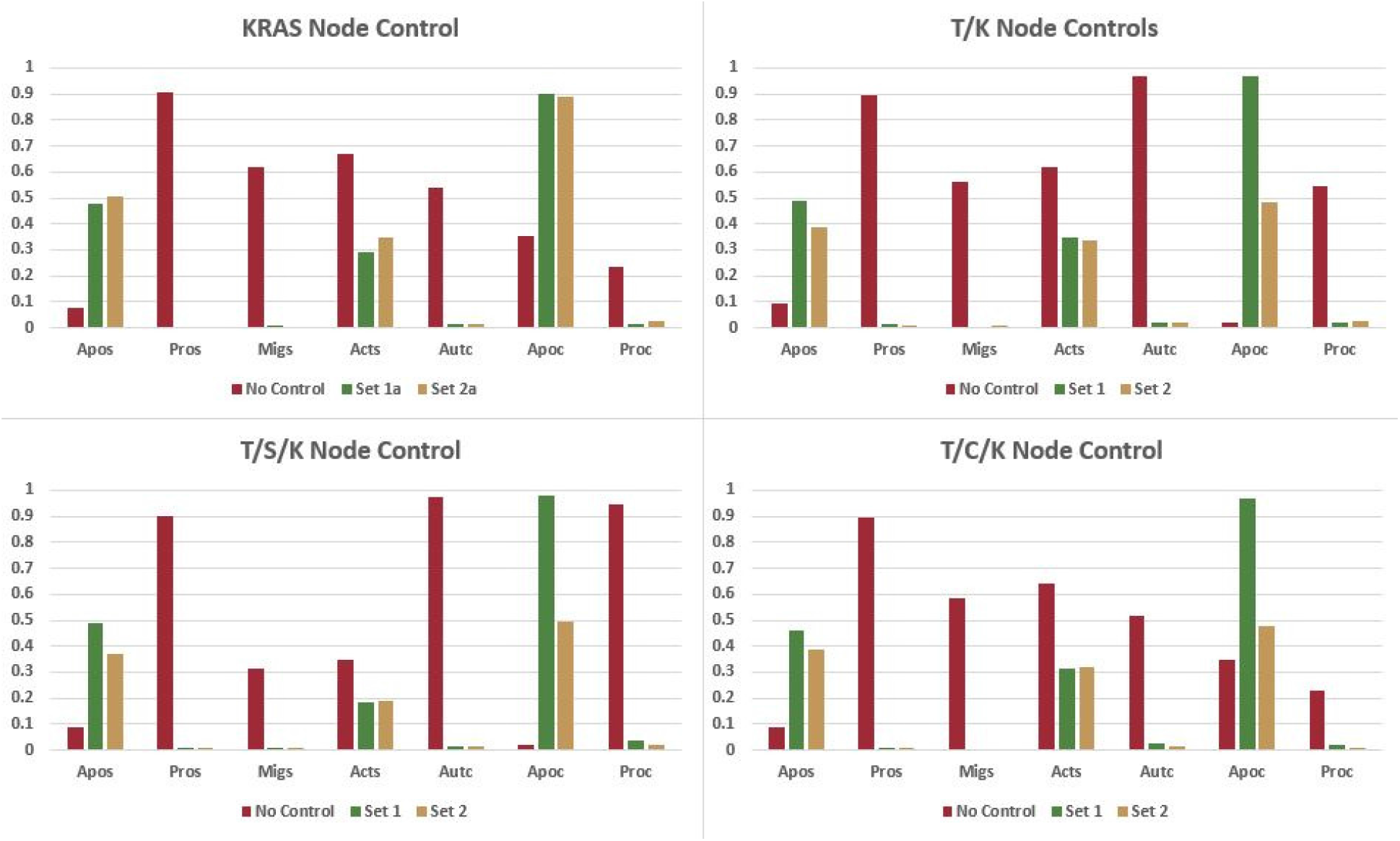
Simulated Node Control. This figure shows a comparison of simulated node control strategies (No Control, Set 1, and Set 2). Expression levels were determined by running 1000 random initializations for 300 time steps (with 1%noise), then recording the average expression at the last time point. Notice that the PSC interventions knockout all PSC phenotypes except activation and apoptosis. We also observe that Set 1 outperforms Set 2 in cases without TP53. See Table 4 for expression levels across all mutation combinations. Set 1: PIK3_c_ / BAX_c_ / ERK_s_/ PIK3_s_ - (Set 1a for KRAS excludes BAX_c_) Set 2: PIP3_c_ / RAF_c_ / ERK_s_ / PIK3_s_ - (Set 2a for KRAS excludes RAF_c_)

**Fig 3.**
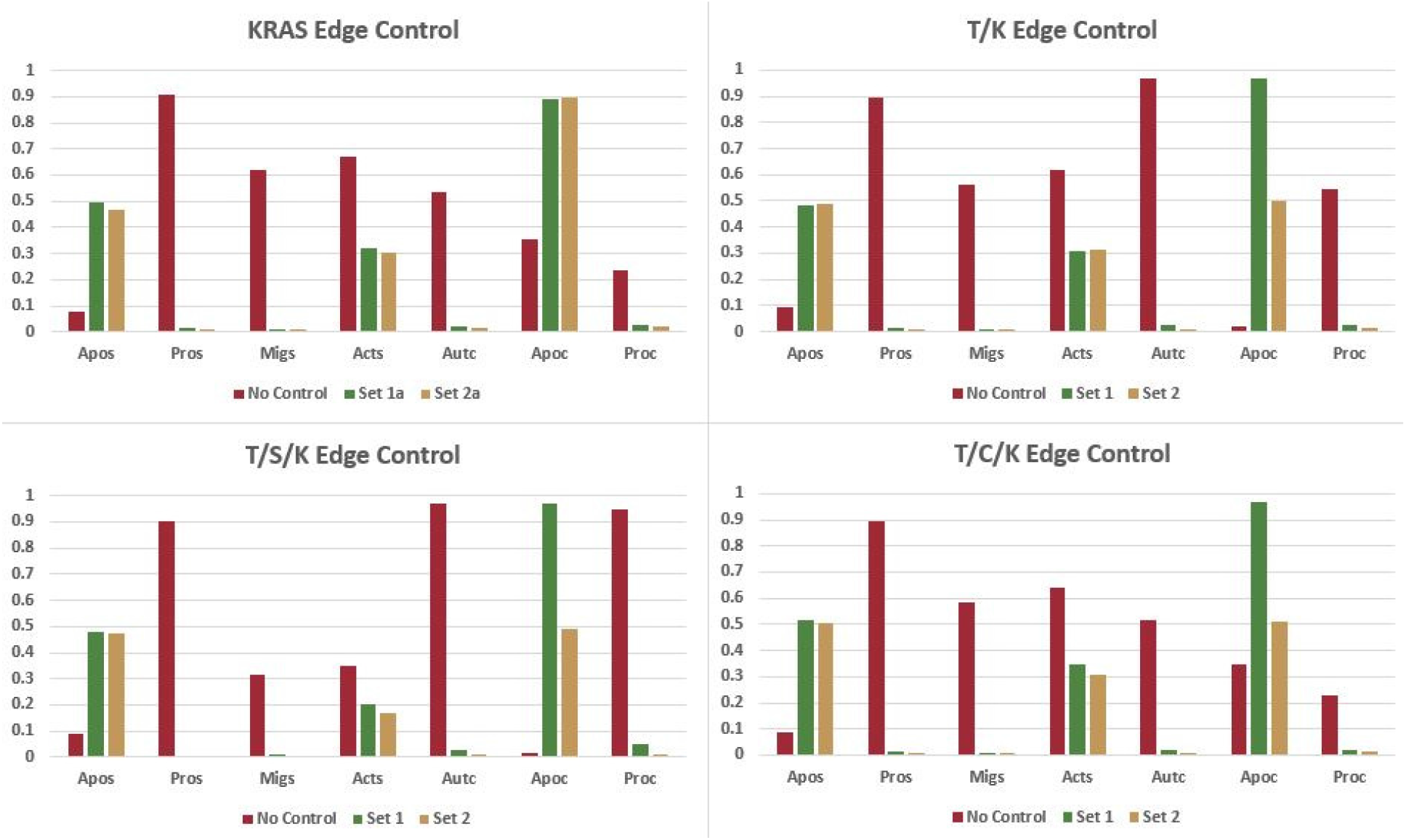
Simulated Edge Control. This figure shows a comparison of simulated edge control strategies (No Control, Set 1, and Set 2). Results from the edge control simulations are nearly the same as those with node control. However, it may be easier to block a signal rather than knockout a gene altogether. Set 1: PIK3_c_→PIP3_c_ / BAX_c_→CASP_c_ /ERK_s_→AP1_s_ / PIK3_s_→PIP3_s_ - (Set 1a for KRAS excludes BAX_c_→CASP_c_) Set 2: PIP3_c_→AKT_c_ / RAF_c_→MEK_c_ / ERK_s_→AP1_s_ / PIK3_s_→PIP3_s_ - (Set 2a for KRAS excludes RAF_c_→MEK_c_)

### 2.2 Gene expression analysis

Given the presence of a particular mutation scenario, certain genes may have more of an adverse response than others (e.g. spikes or dips in expression). The Kruskal-Wallis test was used on the TCGA data set to determine if any median gene expression was significantly different among mutation groups. Using groups with greater than or equal to ten patients, we discovered the results listed in Table 2, which shows the *H* statistic along with the uncorrected p-value. With a significance value of *p* ≤ 0.01, we see that MDM2, BAX, and PIK3CD all show that at least one of the median gene expressions among the combination groups is significantly different. As stated in preceding sections, these three genes proved to be potential controls under the algebraic method. Thus, the statistical analysis performed serves as a validation for model predictions and target discovery.

**Table 2.** Kruskal-Wallis Test. This table shows the results of the Kruskal-Wallis Test for mutation groups with more than 10 patients.

| Gene | H-Value | p-value (uncorrected) | Gene | H-Value | p-value (uncorrected) |
| --- | --- | --- | --- | --- | --- |
| MDM2 | 23.365593 | 0.000287 | RB1 | 7.422619 | 0.191058 |
| BAX | 22.846489 | 0.000361 | CD274 | 7.094339 | 0.213718 |
| PIK3CD | 19.727543 | 0.001406 | FGFR1 | 6.716675 | 0.242579 |
| CCND1 | 13.960565 | 0.015862 | CUX1 | 5.632765 | 0.343607 |
| EGFR | 12.918148 | 0.024158 | TP53 | 5.447952 | 0.363687 |
| BCL2L1 | 10.728332 | 0.057041 | PTEN | 4.414729 | 0.491373 |
| MTOR | 10.315388 | 0.066776 | CCNE1 | 3.958414 | 0.555419 |
| CDKN1A | 9.923672 | 0.077427 | E2F1 | 3.482174 | 0.626087 |
| SMAD4 | 9.470798 | 0.091696 | AKT1 | 3.268281 | 0.658701 |
| RAF1 | 9.384769 | 0.094667 | BECN1 | 2.93573 | 0.709894 |
| TGFBR1 | 8.870499 | 0.114342 | MAPK1 | 2.897151 | 0.715838 |
| MAP2K1 | 8.351973 | 0.137875 | JUN | 2.498611 | 0.776704 |
| KRAS | 7.851126 | 0.164633 | MAPK8 | 2.220305 | 0.817898 |
| PIK3CB | 7.429194 | 0.190627 | NFKB1 | 1.59621 | 0.901707 |

In Fig 4, we give a visual representation of which particular mutation group is different by observing median gene expression levels (white dot within the violin). Note that the violin plot performs smoothing on the edges, which creates negative expression values. The largest difference in median for MDM2 is KRAS, largest difference in median for BAX is KRAS, and largest difference in median for PIK3CD is TP53/CDKN2A/KRAS. To confirm visual results, we performed k-means clustering with *k* = 2. The results in each case showed three medians in one cluster and the fourth median in another, which was classified as the outlier.

**Fig 4.**
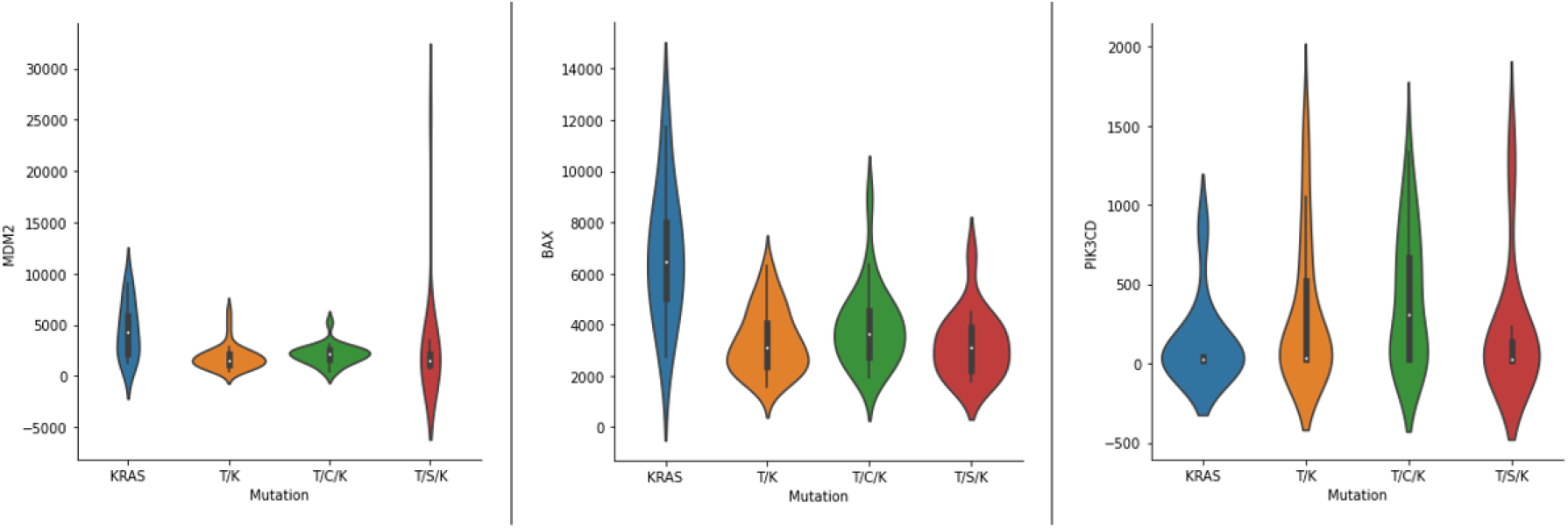
Violin Plots. This figure contains a violin plots of MDM2, BAX, and PIK3CD gene expressions. See Table 2. We found the largest difference in median score by completing *k*-means clustering with *k* = 2. The largest difference in median for MDM2 is KRAS, largest difference in median for BAX is KRAS, and largest difference in median for PIK3CD is T/C/K. Plotting performs smoothing on the edges, which creates negative expression values.

### 2.3 Mutation aggression and control efficacy

Using our model and simulator, we were able to qualitatively generate aggressiveness scores based on attractor probability as well as simulated long-term trajectories, in order to determine if certain mutation combinations are more aggressive than others. We define an attractor as a set of states from which there is no escape as the system evolves, and an attractor with a single state is called a fixed point. The long-term dynamics of BNs will always converge to either a singleton fixed point or a complex attractor. Further details of the scoring weights can be found in Section 4.4. In short, apoptosis was given a negative weight (since it negatively impacts aggression), while positive weights were given to autophagy and proliferation.

We recorded a summary of the probability of landing in a particular attractor (grouped by phenotype in Table 3), and we used a weighted scoring system to achieve the heat maps in Table 4. By inducing mutations, we discovered the creation of new attractors. For example, the non-induced system contained thirty-six total attractors. Yet, combinations such as SMAD4/KRAS yielded thirty-two attractors with only sixteen attractors matching those in the original non-induced system. This can partially be attributed to oscillations being removed by certain mutation inductions. We see that in Table 4, through attractor aggression estimates, combinations of TP53/SMAD4 and TP53/SMAD4/KRAS appear to be the most aggressive, followed closely by TP53 and TP53/KRAS. These simulations had to be run without noise because adding noise makes the system ergodic [1].

**Table 3.** Expression Approximations. This table records the approximate attractor probability for each mutation combination, as well as approximate phenotype expressions. Given 1,000 random initial states, these results show attractor (and trajectory) approximations after 300 time steps (i.e. function updates). This table also records the simulated long-term trajectories for each mutation combination with 1%noise. Then to confirm control efficacy, we ran simulations using control Set 1 (PIK3_c_ / BAX_c_ / ERK_s_ / PIK3_s_) and Set 2 (PIP3_c_ / RAF_c_ / ERK_s_ / PIK3_s_) in the last two sub-tables.

| Uncontrolled Attractor Approximations |  |  |  |  |  |  |  |  |  |  |  |  |  |  |  |  |
| --- | --- | --- | --- | --- | --- | --- | --- | --- | --- | --- | --- | --- | --- | --- | --- | --- |
|  | N.I. | KRAS | TP53 | CycD | SMAD | T-K | C-K | S-K | T-C | T-S | C-S | T-C-K | T-S-K | C-S-K | T-C-S | T-C-S-K |
| Apo<sub>c</sub> | 0 | 0 | 0 | 0 | 0 | 0 | 0 | 0 | 0 | 0 | 0 | 0 | 0 | 0 | 0 | 0 |
| Aut<sub>c</sub> | 0.473 | 0.464 | 0.456 | 0.999 | 0 | 0.422 | 1 | 0 | 1 | 0 | 1 | 1 | 0 | 1 | 1 | 1 |
| Pro<sub>c</sub> | 0.527 | 0.536 | 0 | 0 | 1 | 0 | 0 | 1 | 0 | 0 | 0 | 0 | 0 | 0 | 0 | 0 |
| Aut<sub>c</sub>/Pro<sub>c</sub> | 0 | 0 | 0.544 | 0 | 0 | 0.578 | 0 | 0 | 0 | 0.999 | 0 | 0 | 1 | 0 | 0 | 0 |

| Uncontrolled Trajectory Approximations |  |  |  |  |  |  |  |  |  |  |  |  |  |  |  |  |
| --- | --- | --- | --- | --- | --- | --- | --- | --- | --- | --- | --- | --- | --- | --- | --- | --- |
|  | N.I. | KRAS | TP53 | CycD | SMAD | T-K | C-K | S-K | T-C | T-S | C-S | T-C-K | T-S-K | C-S-K | T-C-S | T-C-S-K |
| Aut<sub>c</sub> | 0.509 | 0.531 | 0.964 | 0.818 | 0.528 | 0.974 | 0.827 | 0.356 | 0.953 | 0.955 | 0.817 | 0.97 | 0.967 | 0.82 | 0.962 | 0.961 |
| Apo<sub>c</sub> | 0.359 | 0.356 | 0.021 | 0.109 | 0.343 | 0.011 | 0.118 | 0.479 | 0.025 | 0.035 | 0.105 | 0.018 | 0.013 | 0.1 | 0.028 | 0.017 |
| Pro<sub>c</sub> | 0.213 | 0.209 | 0.56 | 0.017 | 0.211 | 0.555 | 0.013 | 0.24 | 0.018 | 0.92 | 0.012 | 0.015 | 0.944 | 0.018 | 0.01 | 0.014 |

| Control Set 1 Trajectory Approximations |  |  |  |  |  |  |  |  |  |  |  |  |  |  |  |  |
| --- | --- | --- | --- | --- | --- | --- | --- | --- | --- | --- | --- | --- | --- | --- | --- | --- |
|  | N.I. | KRAS | TP53 | CycD | SMAD | T-K | C-K | S-K | T-C | T-S | C-S | T-C-K | T-S-K | C-S-K | T-C-S | T-C-S-K |
| Aut<sub>c</sub> | 0.016 | 0.028 | 0.012 | 0.021 | 0.014 | 0.019 | 0.021 | 0.022 | 0.02 | 0.016 | 0.019 | 0.023 | 0.014 | 0.018 | 0.02 | 0.022 |
| Apo<sub>c</sub> | 0.978 | 0.962 | 0.979 | 0.968 | 0.973 | 0.97 | 0.965 | 0.963 | 0.97 | 0.966 | 0.979 | 0.969 | 0.981 | 0.975 | 0.975 | 0.969 |
| Pro<sub>c</sub> | 0.025 | 0.029 | 0.021 | 0.016 | 0.027 | 0.021 | 0.015 | 0.028 | 0.015 | 0.026 | 0.018 | 0.019 | 0.035 | 0.012 | 0.012 | 0.019 |

**Table 3. Expression Approximations.**
| Control Set 2 Trajectory Approximations |  |  |  |  |  |  |  |  |  |  |  |  |  |  |  |  |
| --- | --- | --- | --- | --- | --- | --- | --- | --- | --- | --- | --- | --- | --- | --- | --- | --- |
|  | N.I. | KRAS | TP53 | CycD | SMAD | T-K | C-K | S-K | T-C | T-S | C-S | T-C-K | T-S-K | C-S-K | T-C-S | T-C-S-K |
| Aut<sub>c</sub> | 0.008 | 0.019 | 0.016 | 0.019 | 0.015 | 0.019 | 0.006 | 0.014 | 0.013 | 0.019 | 0.017 | 0.013 | 0.015 | 0.026 | 0.02 | 0.01 |
| Apo<sub>c</sub> | 0.946 | 0.934 | 0.503 | 0.942 | 0.932 | 0.485 | 0.934 | 0.926 | 0.517 | 0.497 | 0.933 | 0.475 | 0.496 | 0.925 | 0.49 | 0.475 |
| Pro<sub>c</sub> | 0.01 | 0.02 | 0.013 | 0.016 | 0.013 | 0.024 | 0.015 | 0.021 | 0.014 | 0.025 | 0.013 | 0.011 | 0.019 | 0.015 | 0.022 | 0.014 |

**Table 4.**
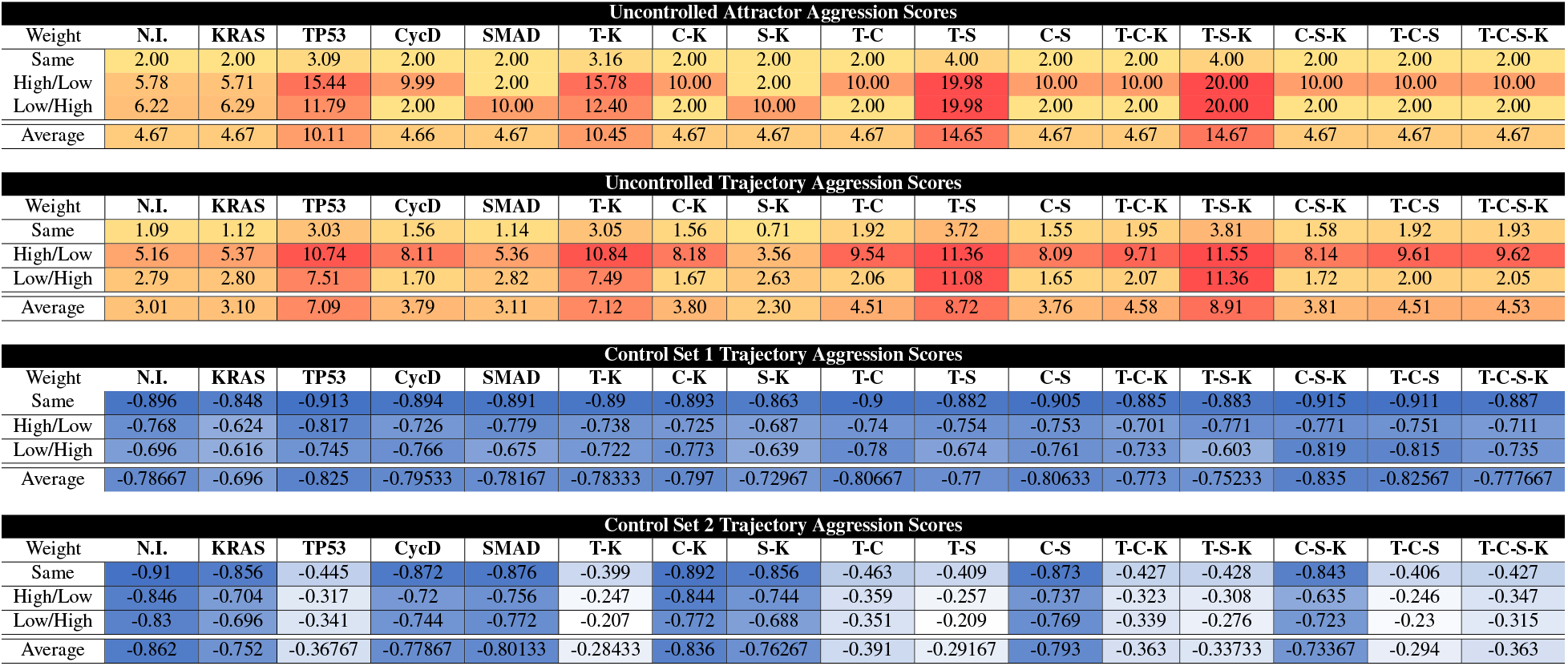
Aggression Scores. This table calculates aggression scores based on the correlating expression levels by applying weights to the attractor (and trajectory) probabilities found in Table 3. Here we include heat maps of aggression scores for each mutation combination (including the case when no mutation was induced (N.I.)), comparing cancer cell autophagy and proliferation while giving a negative weight to apoptosis. Row label “Same” indicates that the same weight was given to both autophagy and proliferation, “High/Low” indicates a high weight for autophagy but a low weight for proliferation, and “Low/High” indicates a low weight for autophagy but a high weight for proliferation. Scaling of the heat map ranges from green (low score) to red (high score) based on the maximum and minimum values. Scaling shades of blue (cold) indicate non-aggressive or negative scores.

In a like manner, we recorded a summary of long-term trajectories and used the same weighted scoring system as before (also in Tables 3 and 4). Trajectory approximations were achieved by running noisy simulations and recording the final average expression levels. Again, we see that combinations of TP53/SMAD4 and TP53/SMAD4/KRAS appear to be the most aggressive, followed closely by TP53 and TP53/KRAS because of high levels of both autophagy and proliferation. Thus, the methods of attractor and trajectory scoring in Table 4 both reveal similar results.

Then to confirm node control efficacy, we ran simulations using control Set 1 (PIK3_c_ / BAX_c_ / ERK_s_ / PIK3_s_) and Set 2 (PIP3_c_ / RAF_c_ / ERK_s_ / PIK3_s_). We created correlating heat maps in Table 4 where scaling blue (i.e. cold) indicates non-aggression. Notice that all controlled aggression scores are negative, and that control Set 1 is most effective across all mutation combinations. However, it appears that Set 2 could have recognizably better impact on the non-induced, KRAS, CDKN2A/KRAS, and SMAD/KRAS mutation scenarios.

### 2.4 Survival curves

Within the TCGA data set, we were able to use survival data to create Kaplan-Meier survival curves for various mutation combinations. Of the 119 patient cohort, 71 were deceased and 48 were alive. In Fig 5 we included the overall survival in addition to the same key combinations used in Section 2.2. This is because the data set does not provide enough points in every combination category to create a sufficient curve (see Table 6 in the *Appendix* for the combination frequencies). The time-to-death variable is recorded in days, and each curve represents the probability of survival for patients in each cohort.

**Fig 5.**
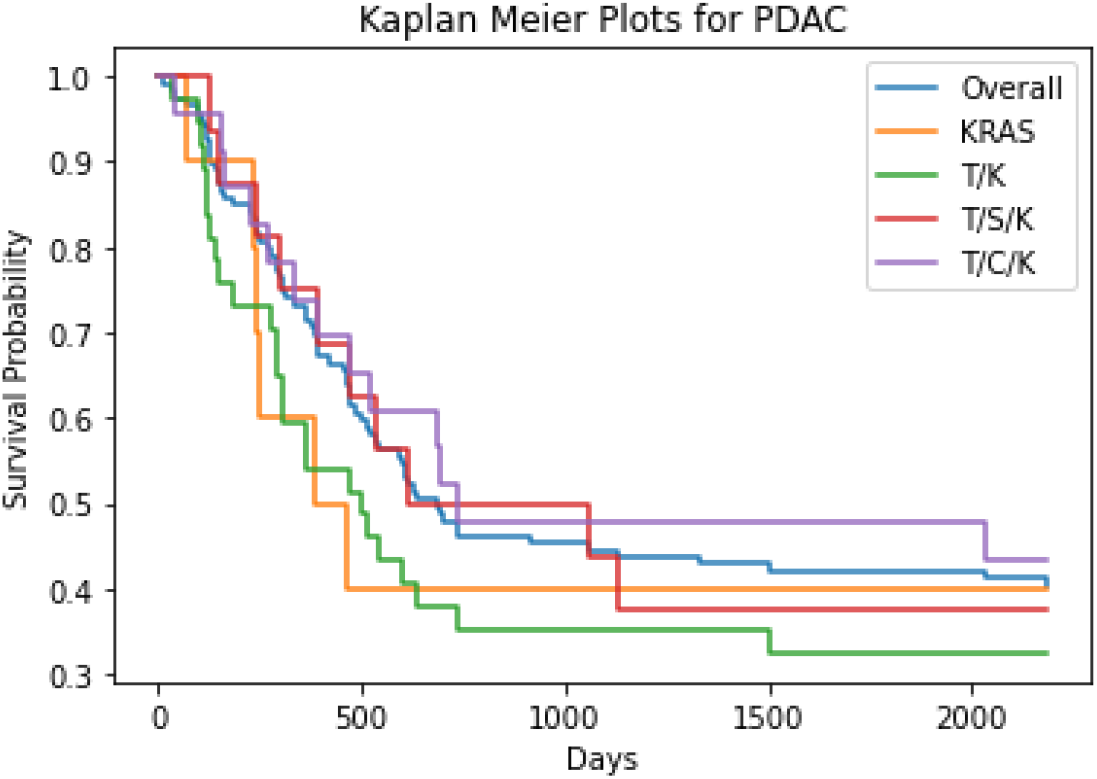
Survival Estimates. This graph shows the Kaplan-Meier survival plots for the key mutation combinations used previously. The time-to-death variable is recorded in days, and each curve represents the probability of survival for patients in each cohort.

**Table 5.** Total mutation frequencies. This table represents the total frequencies of each individual mutation for 119 patients.

|  | Frequency | Percentage |
| --- | --- | --- |
| KRAS | 104 | 87.4% |
| TP53 | 94 | 79.0% |
| SMAD4 | 36 | 30.3% |
| CDKN2A | 34 | 28.6% |

**Table 6.**
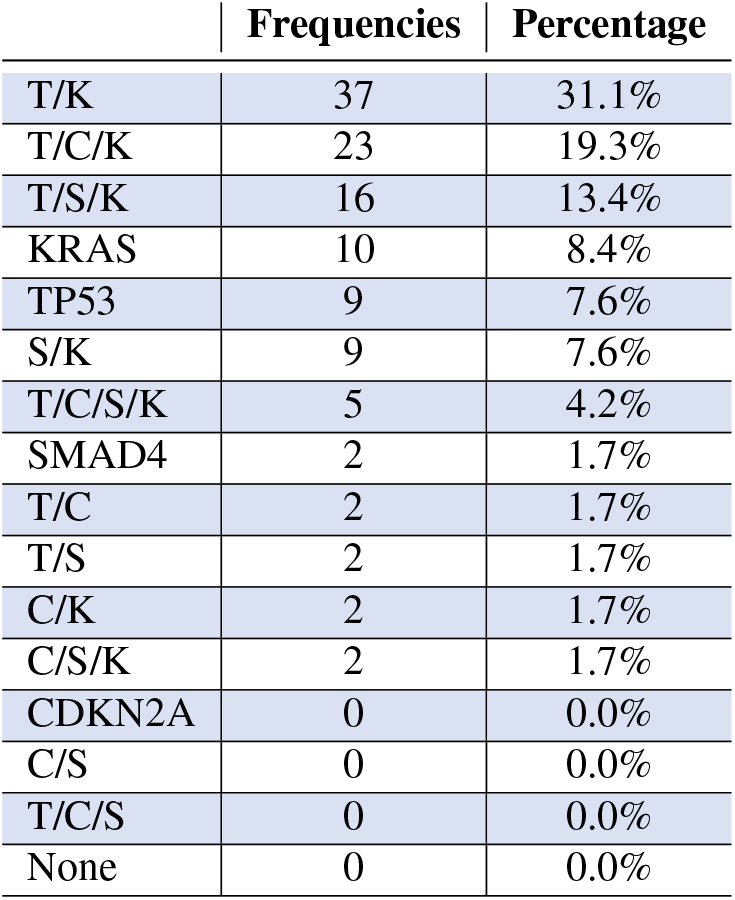
Mutation combination frequencies. This table represents all possible combinations of mutations within our patient sample (119 total patients). Here we abbreviate the mutations as T~TP53, C~CyclinD (CDKN2A), S-SMAD, and K~KRAS.

|  | Frequencies | Percentage |
| --- | --- | --- |
| T/K | 37 | 31.1% |
| T/C/K | 23 | 19.3% |
| T/S/K | 16 | 13.4% |
| KRAS | 10 | 8.4% |
| TP53 | 9 | 7.6% |
| S/K | 9 | 7.6% |
| T/C/S/K | 5 | 4.2% |
| SMAD4 | 2 | 1.7% |
| T/C | 2 | 1.7% |
| T/S | 2 | 1.7% |
| C/K | 2 | 1.7% |
| C/S/K | 2 | 1.7% |
| CDKN2A | 0 | 0.0% |
| C/S | 0 | 0.0% |
| T/C/S | 0 | 0.0% |
| None | 0 | 0.0% |

**Table 7.**
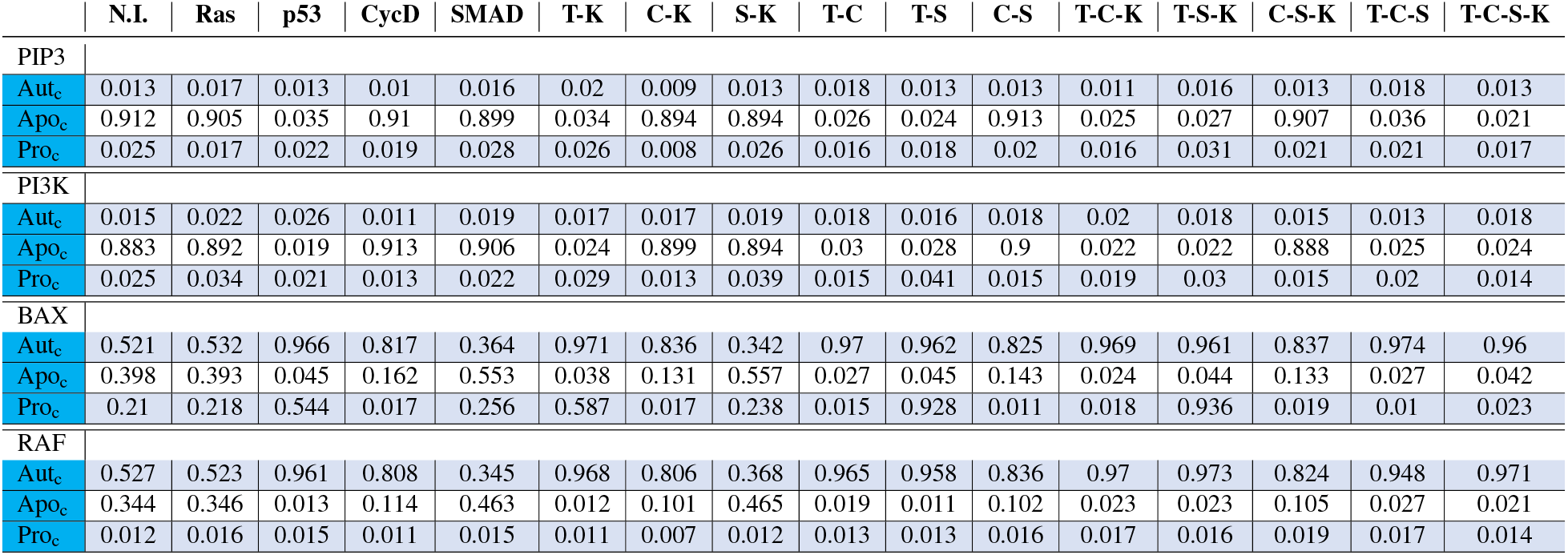
Singleton PCC node control. This figure shows the long term trajectories of using singleton node controls. Expression levels were determined by running 1000 random initializations for 300 time steps (with 1%noise), then recording the average expression at the last time point.

|  | N.I. | Ras | p53 | CycD | SMAD | T-K | C-K | S-K | T-C | T-S | C-S | T-C-K | T-S-K | C-S-K | T-C-S | T-C-S-K |
| --- | --- | --- | --- | --- | --- | --- | --- | --- | --- | --- | --- | --- | --- | --- | --- | --- |
| PIP3 |  |  |  |  |  |  |  |  |  |  |  |  |  |  |  |  |
| Aut<sub>c</sub> | 0.013 | 0.017 | 0.013 | 0.01 | 0.016 | 0.02 | 0.009 | 0.013 | 0.018 | 0.013 | 0.013 | 0.011 | 0.016 | 0.013 | 0.018 | 0.013 |
| Apo<sub>c</sub> | 0.912 | 0.905 | 0.035 | 0.91 | 0.899 | 0.034 | 0.894 | 0.894 | 0.026 | 0.024 | 0.913 | 0.025 | 0.027 | 0.907 | 0.036 | 0.021 |
| Pro<sub>c</sub> | 0.025 | 0.017 | 0.022 | 0.019 | 0.028 | 0.026 | 0.008 | 0.026 | 0.016 | 0.018 | 0.02 | 0.016 | 0.031 | 0.021 | 0.021 | 0.017 |
| PI3K |  |  |  |  |  |  |  |  |  |  |  |  |  |  |  |  |
| Aut<sub>c</sub> | 0.015 | 0.022 | 0.026 | 0.011 | 0.019 | 0.017 | 0.017 | 0.019 | 0.018 | 0.016 | 0.018 | 0.02 | 0.018 | 0.015 | 0.013 | 0.018 |
| Apo<sub>c</sub> | 0.883 | 0.892 | 0.019 | 0.913 | 0.906 | 0.024 | 0.899 | 0.894 | 0.03 | 0.028 | 0.9 | 0.022 | 0.022 | 0.888 | 0.025 | 0.024 |
| Pro<sub>c</sub> | 0.025 | 0.034 | 0.021 | 0.013 | 0.022 | 0.029 | 0.013 | 0.039 | 0.015 | 0.041 | 0.015 | 0.019 | 0.03 | 0.015 | 0.02 | 0.014 |
| BAX |  |  |  |  |  |  |  |  |  |  |  |  |  |  |  |  |
| Aut<sub>c</sub> | 0.521 | 0.532 | 0.966 | 0.817 | 0.364 | 0.971 | 0.836 | 0.342 | 0.97 | 0.962 | 0.825 | 0.969 | 0.961 | 0.837 | 0.974 | 0.96 |
| Apo<sub>c</sub> | 0.398 | 0.393 | 0.045 | 0.162 | 0.553 | 0.038 | 0.131 | 0.557 | 0.027 | 0.045 | 0.143 | 0.024 | 0.044 | 0.133 | 0.027 | 0.042 |
| Pro<sub>c</sub> | 0.21 | 0.218 | 0.544 | 0.017 | 0.256 | 0.587 | 0.017 | 0.238 | 0.015 | 0.928 | 0.011 | 0.018 | 0.936 | 0.019 | 0.01 | 0.023 |
| RAF |  |  |  |  |  |  |  |  |  |  |  |  |  |  |  |  |
| Aut<sub>c</sub> | 0.527 | 0.523 | 0.961 | 0.808 | 0.345 | 0.968 | 0.806 | 0.368 | 0.965 | 0.958 | 0.836 | 0.97 | 0.973 | 0.824 | 0.948 | 0.971 |
| Apo<sub>c</sub> | 0.344 | 0.346 | 0.013 | 0.114 | 0.463 | 0.012 | 0.101 | 0.465 | 0.019 | 0.011 | 0.102 | 0.023 | 0.023 | 0.105 | 0.027 | 0.021 |
| Pro<sub>c</sub> | 0.012 | 0.016 | 0.015 | 0.011 | 0.015 | 0.011 | 0.007 | 0.012 | 0.013 | 0.013 | 0.016 | 0.017 | 0.016 | 0.019 | 0.017 | 0.014 |

We see that the combination TP53/KRAS seems to have the worst survival rate among patients in this cohort, followed by the combination TP53/SMAD4/KRAS. The results in Section 2.3 (Table 4) suggested that, via simulation, TP53/SMAD4 and TP53/SMAD4/KRAS are the most aggressive, followed closely by TP53 and TP53/KRAS. This was consistent with the results from our model for the cases with sufficient data points among cohorts, as we were only able to create survival curves for combinations with greater than or equal to ten patients (see Table 6).

## 3 Discussion and conclusion

We have built a model to study the impact of inducing four common mutations: KRAS, TP53, SMAD4, and CDKN2A. Sets of control targets were discovered through computational algebra and are effective among common mutations, including PIP3c, RAFc, PIK3c, BAXc, ERKs, and PIK3s. After implementing the various mutation combinations, we used our simulator to derive aggressiveness scores based on simulated attractor probability and long-term trajectories. Results indicated that certain mutation combinations could potentially be more aggressive than others, in the sense that they are more difficult to treat or that the patient might see a faster decline. Statistical analysis was also performed on clinical gene expression data to help corroborate the results of our model.

Recent efforts have shown promise in controlling KRAS, which was previously believed to be uncontrollable [29]. However, within our simulator, inducing a mutation will not allow that particular gene to be controlled. For example, applying KRAS control on the TP53/KRAS mutation would result in the expression levels seen for TP53 because the mutation induction would merely be reversed. Moreover, we recognize slight variance between model simulations and clinical gene expression data. One possible cause is that our model is multicellular, while the clinical data is single cell. Even though the model considers the microenvironment, BNs can be an oversimplification of an extremely complex system. Further, the small data set also presented a challenge because we were unable to validate the entirety of our model predictions. Nonetheless, the overall results appear to be consistent with the data.

The Kruskal-Wallis test told us that at least one of the medians, among genes grouped by mutation combination, is different than the others. Therefore, we would expect the simulated expression levels to indicate the same results as Fig 4. While we do see similarities, we note some discrepancies in the simulation results shown in Fig 6 in the *Appendix*. High levels of PIK3 are observed because the topology of the network yields a feedback loop through the cancer cell MAPK pathway, which leads to constant expression of PIK3 if KRAS is mutated (as is the case in every combination considered here, see Fig 1). Additionally, downstream signalling through cancer cell TP53/…/AKT/NFκB leads to feedback from the stellate cell MAPK pathway to overexpress bFGF, which results in the same outcome. Lastly, the induced TP53/CDKN2A/KRAS mutation causes constant expression of MDM2, which would otherwise be oscillatory and show lower expression levels.

**Fig 6.**
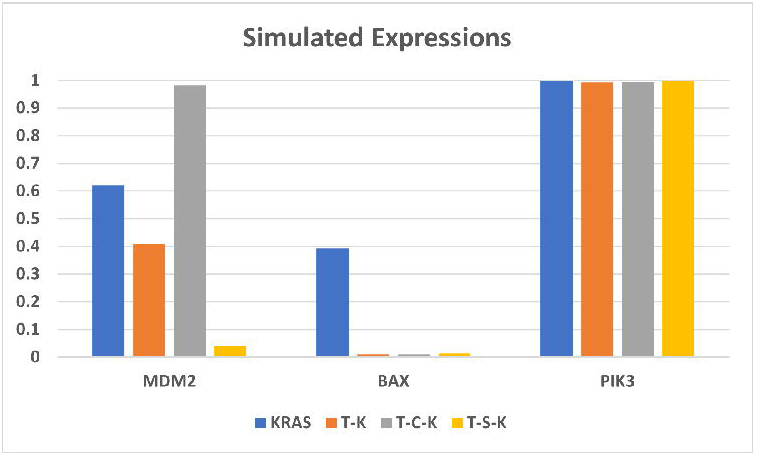
Simulation gene expressions. This figure shows the simulated gene expressions for significant genes found in Table 2 with 1%noise. As with other simulations, levels were approximated by running 1000 random initializations for 300 time steps, then recording the average frequency at the final time step. The chart is grouped by genes, showing each mutations impact on simulated expression.

In a BN the magnitude of the state space for *n* genes is 2^*n*^, which means that an increase of network size will exponentially increase the computational burden for its analysis. Therefore, brute-force methods that can be used on small systems are not sufficient. When such a situation arises, there are reduction techniques available to reduce the computational load, while maintaining important system dynamics. Notably, methods of stable motifs and feedback vertex set (FVS) require apoptotic attractors to determine control targets [27]. However, we were able to avoid magnitude issues because inducing KRAS, TP53/KRAS, TP53/SMAD4/KRAS, and TP53/CDKN2A/KRAS mutations created strictly diseased attractors (autophagy and apoptosis). Yet, there are certain combinations that maintained healthy attractors (apoptosis), such as CDKN2A, TP53, TP53/CDKN2A, TP53/SMAD4, CDKN2A/SMAD4, and TP53/CDKN2A/SMAD4, and one could use stable motifs [41] or FVS [42] to drive the systems to their respective apoptotic attractors.

In Table 7 (see *Appendix*), we show the simulated expressions for individual node knockout in order to verify that specific mutation combinations may require a combination of drugs. This is consistent with findings in [34], where they tested the efficacy of drugs on pathways similar to those found in our model, and they showed evidence that particular drugs may operate better in combination rather than individually. Even so, Table 4 indicates heterogeneity in controls for multiple mutational states. We observed that Set 1 (PIK3_c_ / BAX_c_ / ERK_s_ / PIK3_s_) shows the best overall success in aggression depletion, whereas Set 2 (PIP3_c_ / RAF_c_ / ERK_s_ / PIK3_s_) shows more effective control in specific scenarios. To that end, the computational algebra method provides a substantial list of control options to parse, and those presented herein are the most effective that we found. The details of how we selected the effective node and edge controls are given in Section 4.3.

## 4 Methods

All computations were performed on an Intel(R) Core i7-10750H CPU @ 2.60GHz, 2592 Mhz, 6 Core(s), 12 Logical Processor(s) with 16 GB of RAM and a 64-bit operating system.

### 4.1 Stochastic discrete dynamical systems

Synchronous updating schedules produce deterministic dynamics, where all nodes are updated simultaneously (i.e. in sync). The stochastic discrete dynamical systems (SDDS) framework developed by [22] incorporates Markov chain tools to study long-term dynamics of Boolean networks. By definition, an SDDS of the variables (x_1_;*x_2_*,…, x_n_) is a collection of *n* triples denoted 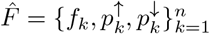, where for *k* = 1,…,*n*,

- f_k_: {0,1}^n^ → {0,1}is the update function for *x_k_*
- 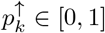 is the activation propensity
- 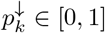 is the deactivation propensity

Consider the state-space *S*, consisting of all possible states of the system. If *x* = (*x*_1_,…, *x_n_*) ∈ *S* and *y* = (*y*_1_,…, *y_n_*) ∈ *S*, then the probability of transitioning from *x* to *y* is

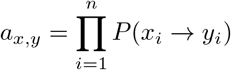

where

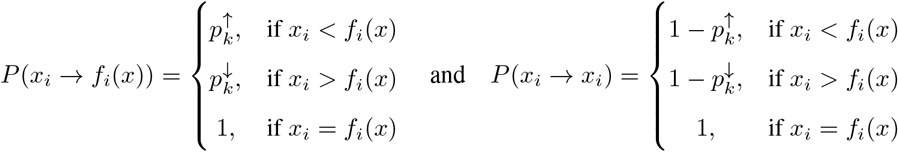

Then *P*(*x_i_* → *y_i_*) = 0 for any *y_i_* ∈ {*x_i_*, *f_i_*(*x*)}. When propensities are set to *p* = 1, we have a traditional BN [27].

With this framework, we built a simulator that takes random initial states as inputs and then tracks the trajectory of each node through time. Long-term phenotype expression probabilities can then be estimated, as well as network dynamics with (and without) controls.

### 4.2 Model creation

In [27], we detail the construction of our multicellular pancreatic cancer model. To encode signaling pathways, we used a Boolean network where each node is either ON or OFF (1 or 0). In other words, the node is either being activated or deactivated. We then can determine the state of the entire system by the node-wise states at current time. Node activation was written with “OR” statements, while deactivation was written as “AND NOT” statements.

### 4.3 Control method based on computational algebra

The method based on computational algebra described in [24] seeks two types of controls: nodes and edges. These can be achieved biologically by blocking effects of the products of genes associated with nodes, or targeting specific gene communications. Let the function 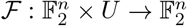 (where 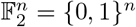 denote a Boolean network with control, where *U* is a set of all possible controls. Then, for some *u* ∈ *U*, the new system dynamics are 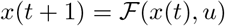. That is, each coordinate *u_i,j_* ∈ *u* encodes the control of edges as follows: consider the edge from nodes *x_i_* → *x_j_* in a given diagram. By construction we have

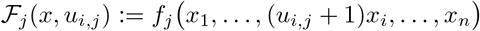

which gives

- Inactive control:

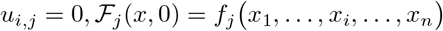
- Active control (edge deletion):

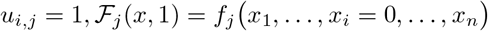

The definition of edge control can therefore be applied to many edges, obtaining 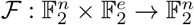 where e is the number of edges in the diagram. Next, we consider control of node *x_i_* from a given diagram. By construction

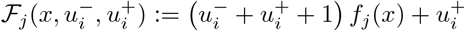

which yields

- Inactive control:

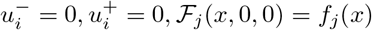
- Node *x_i_* deletion:

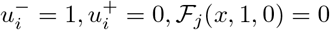
- Node *x_i_* expression:

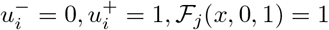
- Negated function value (irrelevant for control):

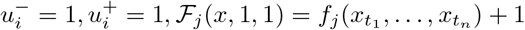

Using these definitions, we can achieve three types of objectives. Let 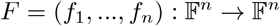 where 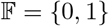 and *μ* = {*μ*_1_,…, *μ_n_*} is a set of controls. Then we can:

- *Generate new attractors*. If *y* is a desirable state (ex. apoptosis), but it is not currently an attractor, we find a set *μ* so that we can solve

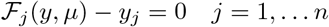
- *Block transitions or remove attractors*. If y is an undesirable attractor (ex. proliferation), we want to find a set *μ* so that 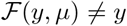. In general, we can use this framework to avoid transitions between states (say *y* → *z*) so that 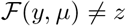. So we can solve

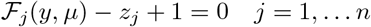
- *Block regions*. If a particular value of *y*, say *y_k_* = *a*, triggers an undesirable pathway, then we need all attractors to satisfy *y_k_* = *a*. So we find a set *μ* so that the following system has no solution

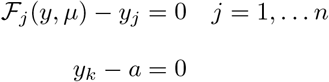

Notably, the Boolean functions F must be written as polynomials to complete the control search by computing the Grobner basis of the ideal associated with the given objective. For example, if we generate new attractors, we find the Grobner basis for

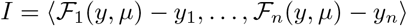

Therefore, we can determine all controls that solve the system of equations and detect combinatorial actions for the given model using the Macaulay2 software [15, 24]. However, the list of controls are merely theoretical controls and must be parsed to find tangible targets (or combinations of targets). The parsing process can include brute-force testing of all controls, knowledge of the regulatory network topology, knowledge of literature pertaining to particular controls, or a mixture of various techniques [24]. Some controls may not be biologically achievable, others may be insufficient if applied independently, while some simply do not perform as desired. To determine the efficacy of controls, we compare uncontrolled simulations with the appropriate target control simulations. Inducing mutations will result in high levels of diseased phenotypes. Thus, a good control will produce low disease levels and high health levels (apoptosis in our case).

### 4.4 Aggression scores

We derived aggressiveness scores for each mutation combination using attractor probability as well as long-term trajectory approximations. Stable motifs [41] were used to derive attractors associated with each mutation combination (see [27, 41] for more details of this method), and they were grouped according to phenotype. We then ran simulations and recorded an approximate probability of landing in a given attractor (i.e. achieving a specific phenotype) in Table 3. For example, the non-induced system contained thirty-six attractors, of which we have four associated with apoptosis, eight with autophagy, and twenty-four with proliferation. After using our simulator with 1000 random initializations, 473 converged to attractors classifying autophagy and 527 converged to attractors classifying proliferation. Here, we do not use noisy simulations because noise makes the system ergodic. On the other hand, we derived long-term expressions (see Table 3) in the same manner but with 1%noise. We see that the non-induced (N.I.) system showed levels of 51%autophagy, 36%apoptosis, and 21%proliferation.

The heat maps in Table 4 are sorted with column-wise mutation groups and used to compare cancer cell autophagy and proliferation while giving a negative weight (*ω* = −1) to apoptosis. The row label “Same” indicates that the same weight was given to both autophagy and proliferation (used value *ω* = 2 for both), “High/Low” indicates a high weight for autophagy (*ω* = 10) but a low weight for proliferation (*ω* = 2), and “Low/High” indicates a low weight for autophagy (*ω* = 2) but a high weight for proliferation (*ω* = 10). Scaling of the heat map ranges green (low score) to red (high score) based on the maximum and minimum values in each table. However, blue shading (i.e. cold) indicates a negative score, which is interpreted as successful depletion of aggression.

Lastly, we justify the positive weight given to autophagy, which is a natural process where cells heal themselves.

The cell will break down any damaged or unnecessary components, and it will reallocate the nutrients from these processes to those that are essential. However, studies have shown that autophagy is required for pancreatic tumor growth [39]. Autophagy can help tumors overcome conditions such as hypoxia and nutrient deprivation. Within tumors, cells can exist under hypoxic conditions. If activated autophagy is then suppressed by deletion of Beclin 1, studies have shown increased cell death. It has also been observed that autophagy is increased in KRAS mutated cells, and aids in survival of the cancer cells while experiencing nutrient starvation. Further, animal studies have shown that autophagy contributes to tumor-cell survival by enhancing stress tolerance and supplying nutrients to meet the metabolic demands of tumors. Then once suppression of autophagy occured, there was an observance of tumor-cell death [40].

### 4.5 Statistical testing

We used the Kruskal-Wallis H-test to find if there was a difference in median gene expressions based on mutation combination. The Kruskal-Wallis test is a nonparametric method with the null hypothesis that the mean ranks of the groups are the same. This test does not assume a normal distribution of the underlying data [38]. All data is ranked from smallest (1) to largest (N), then the sums of ranks in each subgroup are added up, and the probability is calculated.
 The statistic *H* is calculated as follows:

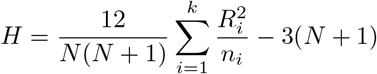

where *N* is the total number of observations across all groups, k is the number of groups, *n_i_* is the number of observations in the ith group, and *R_i_* is the total sum of ranks in the *i*th group. The value of *H* is then tested against the chi-square distribution for *k* – 1 degrees of freedom. If there are tied ranks, then a correction is used [16]. Software from [31] was used in Python3 to encode our statistical testing.

### 4.6 Survival curves

Kaplan-Meier curves and estimates of survival data are a popular way of dealing with various time-to-event data. Time-to-event is a duration variable, where each subject has an initial reading and an end reading anywhere along the timeline of the study. Therefore, time is serial (clinical-course) rather than secular (calendar). For example, initial data is collected when a subject is enrolled into a study or when treatment begins, and it ends when the study has concluded or when the subject is censored (i.e. does not finish) [17]. Survival at the next time step can be calculated by

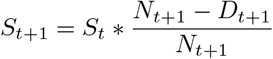

where *S_t_* is the previous survival probability, *N*_*t*+1_ is the number at risk, and *D*_*t*+1_ is the number of deaths [19].

Here, the time-to-event parameter is days-to-death. The variable is representative of the number of days from which the patient was initially screened to the day the patient deceased. In the case where a patient was still alive, we replaced the designated “[Not Applicable]” time with the maximum survival time point. In the same manner as before, we grouped patients by mutation combination in order to compare survival estimates, which were encoded in Python3 using appropriate packages [10].

## Acknowledgements

The results shown here are in whole or part based upon data hosted by the Institute for Systems Biology Cancer Gateway in the Cloud (ISB-CGC) platform. ISB-CGC has been funded in whole or in part with Federal funds from the National Cancer Institute, National Institutes of Health, Task Order No. 17X053 under Contract No. HHSN261200800001E. DM received funding from the Simons Foundation (850896).

## 5 Appendix

All data and code used for running simulations, statistical analysis, and plotting is available on a GitHub repository at https://github.com/drplaugher/PCC_Mutations.

### 5.1 Aggregate statistics

Using the TCGA pancreatic cancer database, we were able to categorize 119 patients according to mutation combination. In Table 6, we see the counts for each combination listed according to prevalence. We also tracked the counts of the individual mutation occurrence, regardless of combination (see Table 5).

### 5.2 Supplementary graphs and tables

